# *Mycobacteria tuberculosis* PPE36 modulates host inflammation by promoting E3 ligase Smurf1-mediated MyD88 degradation

**DOI:** 10.1101/2021.01.07.425701

**Authors:** Zhangli Peng, Yan Yue, Sidong Xiong

## Abstract

*Mycobacterium tuberculosis* (Mtb) PPE36, a cell-wall associated protein is highly specific and conserved for the Mtb complex group. Although it has been proven essential for iron utilization, little is known about the role of PPE36 in regulating host immune responses. Here we exhibited that PPE36 preferentially enriched in Mtb virulent strains, and could efficiently inhibit host inflammatory responses and increase bacterial loads both in mycobacterium-infected macrophages and mice. In exploring the underlying mechanisms, we found that PPE36 could robustly inhibit the activation of inflammatory NF-κB and MAPK (ERK, p38 and JNK) pathways by promoting E3 ligase Smurf1-mediated ubiquitination and proteasomal degradation of MyD88 protein. Our research revealed a previously unknown function of PPE36 on modulating host immune responses, and provided some clues to the development of novel tuberculosis treatment strategies based on immune regulation.

**Author Summary:** Mycobacterium tuberculosis (Mtb) has developed diverse immune evasion strategies to successfully establish infection in host. Identifying the important Mtb immune regulatory proteins and elucidating the underlying mechanisms are critical for tuberculosis control. Here we demonstrated that PPE36, a Mtb cell-wall associated protein, was predominantly enriched in virulent mycobacterial strains, and obviously inhibited inflammatory responses and facilitated bacterial survival in infected macrophages. Compared with the wild-type BCG, BCG lacking PPE36 (BCGΔPPE36) induced more inflammation, lower bacterial loads as well as the improved histopathological changes in the lungs of infected mice. We further found that PPE36 significantly reduced host MyD88 abundance, and inhibited the activation of subsequunt inflammatory NF-κB and MAPK pathways. In addition, this direct inhibition effect of PPE36 on MyD88 was mediated by the promoted E3 ligase Smurf1 ubiquitin -protesome pathway. This study identified PPE36 as a immune regulatory protein of Mtb, and showed it played an important role in the Mtb immune evasion.

## Introduction

Tuberculosis is a global health emergency with about 10 million new TB cases and 1.6 million death in 2017 [1]. Although tuberculosis deaths in many countries appeared rapid declined over past decades, the annual incidence rate decline was only 1.5-2.0% during 2000-2015, less than half of the rate (4-5%) needed to achieve the milestone case reductions and end tuberculosis [1]. In addition, inoculation of the only widely used TB vaccine bacillus Calmette-Guérin (BCG) could not be effective for all populations, therefore the global burden of tuberculosis remains substantial.

As an extremely successful pathogen, *Mycobacterium tuberculosis* (Mtb) has co-evolved with human being for thousands of years, and developed diverse mechanisms to establish latent, progressive or persistent infections in fully immunocompetent hosts. Host macrophages are the initial and primary target of Mtb. In there, they resident and persist by a variety of immune evasion strategies, including hiding from PRR recognition [2], preventing T cell responses by down-regulating MHC molecules [3], as well as evading macrophage killing systems [3] and so on. In addition, the genomic plasticity might also be partially responsible for the antigenic variations of diverse Mtb stains and the variable protective efficacies of diverse BCG strains [4].

Actually, some important comparative analyses have showed us the panoramic views of genome variations among BCG vaccine strains [4, 5]. Kanglin Wan et al [6] showed 43 common proteins between 13 BCG strains. Most of them belonged to PE and PPE protein families with unknown functions. PE and PPE are two families of glycine-rich proteins with a repetitive structure; their nomination was derived from the conserved N-terminal Pro-Glu (PE) and Pro-Pro-Glu (PPE) motifs. In addition to representing the principal source of antigenic variation, more and more PE/PPE proteins exhibited abilities to modulate host immune responses [7].

In this study, we focused on exploring the potential role of PPE36 in the interaction between Mtb and host macrophages. PPE36 is a 27-kDa cell-wall associated protein, and limitedly expressed among the Mtb complex group (including notably Mycobacterium tuberculosis, *Mycobacterium africanum*, *Mycobacterium bovis*) [8–10]. A recent study revealed that PPE36 was essential for the iron acquisition from heme by Mtb [11]. Interestingly, there seemed to be a paradox about PPE36. On one hand, it is dispensable for mycobacterial survival and growth [12], on the other hand it is ubiquitous in various bacterial strains [6, 13]. We hypothesized that PPE36 might have other functions beneficial for Mtb survival and/or infection. In fact, several cell-membrane associated PPE proteins (such as PPE18, PPE34,) have shown to interact with host immune systems, like binding TLR2 [14, 15], decreasing MHC molecules [14, 15]. More importantly, PPE37, another PPE membrane facilitating iron utilization by Mtb just like PPE36, was found to be able to decrease pro-inflammatory cytokine production in the Mtb-infected macrophages [16, 17]. All these works further prompted us to explore the possible influence of PPE36 on macrophage immune responses against Mtb infection.

In this study, we found that PPE36 could promote the ubiquitination and degradation of host MyD88 protein by facilitating the interaction of MyD88 with the E3 ligase Smurf1, and then suppressed the subsequent NF-κB as well as MAPK (ERK, p38 and JNK) pathways, which led to the dampened host inflammatory responses and increased intracellular bacterial loads. Our findings revealed a previously unknown immune-modulation function of PPE36 protein. It will help us better understand the complex immune-evasion mechanisms of Mtb, and may also provide some clues to the development of novel preventive and therapeutic strategies based on PPE proteins.

## Results

### PPE36 was predominantly enriched in virulent but not attenuated mycobacteria

The PPE36 expression in BCG strain, Mtb H37Rv strain or 20 Mtb clinical isolates were detected by the absolute quantification real-time PCR assays. It was found that PPE36 expression was much more enriched in the virulent H37Rv strain compared with the attenuated BCG strain, and its expression was further significantly increased in almost 20 clinical isolates compared with H37Rv (Fig 1A), indicating that similar like other PPE family members [18], PPE36 also preferentially expressed in the virulent mycobacteria.

**Fig 1.**
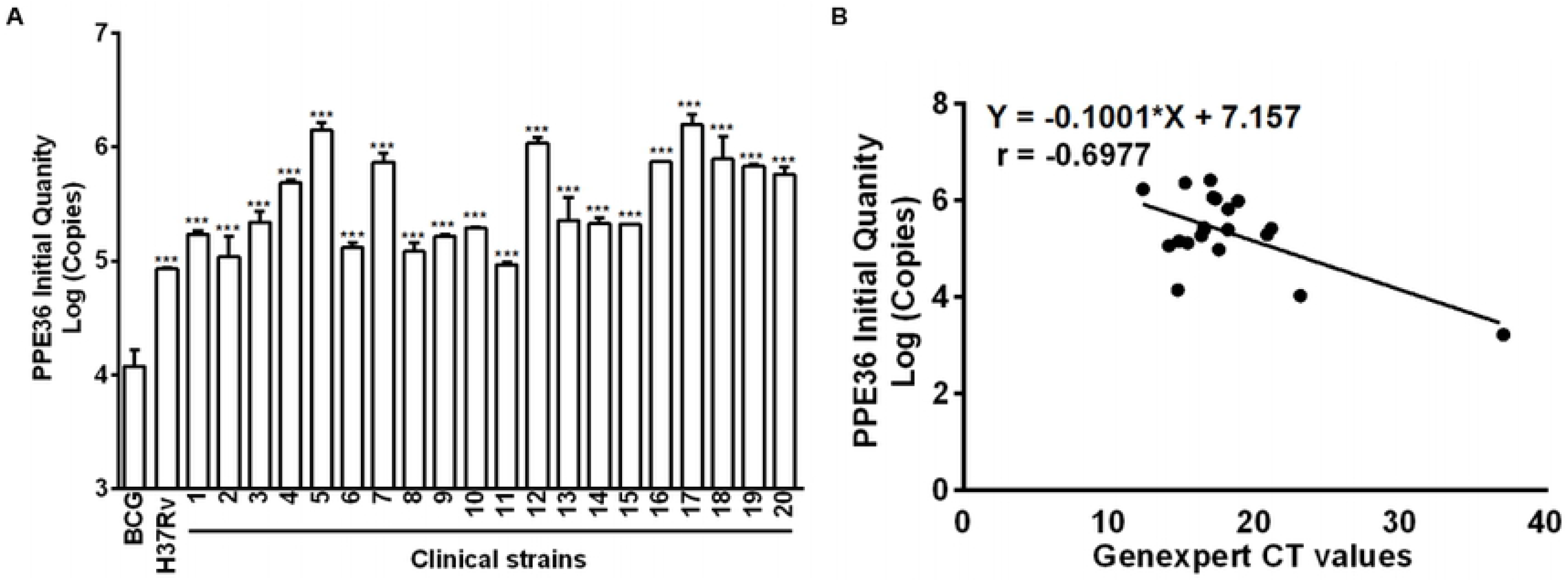
PPE36 expression was predominantly enriched in virulent mycobacteria. (A) PPE36 expression in BCG, H37Rv and clinical isolates (n=20) was quantified by real-time PCR. (B) Correlation of PPE36 expression in clinical isolates with the bacterial loads in clinical samples. Individual experiments were conducted three times with similar results. *P < 0.05, **P < 0.01, ***P <0.001.

Next, we explored the correlation of bacterial PPE36 gene abundance with the bacterial loads in clinical sputum and BALF samples. Herein we applied GeneXpert PCR assays to detect sputum or BALF Mtb, and used Ct value to indirectly reflect the sputum bacterial loads. The more bacterial load in sputum, the lower the GeneXpert CT value will be, because it is easier to be detected. Of interest, we found that PPE36 gene abundance was significantly and negatively correlated with the GeneXpert CT values (r=-0.6977) (Fig 1B), which meant that PPE36 gene abundance was closely and positively correlated with the bacterial loads in the clinical samples. These data suggested that PPE36 might play an important role in Mtb infection and tuberculosis disease process.

### PPE36 suppressed macrophage inflammatory cytokine production

To test whether PPE36 influenced the mycobacterium infection, murine BMDMs were infected with WT or PPE36-forcing expressed *M.smegmatis* (PPE36-*M.smegmatis*), and the levels of inflammatory cytokines were measured by ELISA assays. It was showed that the levels of TNF-α, IL-6 and IL-1β in PPE36-*M.smegmatis* infected cells were significantly decreased compared with those of WT *M.smegmatis* infected cells (TNF-α: 490 vs 400 pg/ml; IL-6: 670 vs 460 pg/ml; IL-1β: 1100 vs 690 pg/ml, Fig 2A). In line with the weakened macrophage inflammation, the intracellular bacterial load also significantly increased in PPE36-*M.smegmatis* infected group and achieved up to 800 CFU/ml, which was about twice that of the control *M.smegmatis* infected group (Fig 2B).

**Fig 2.**
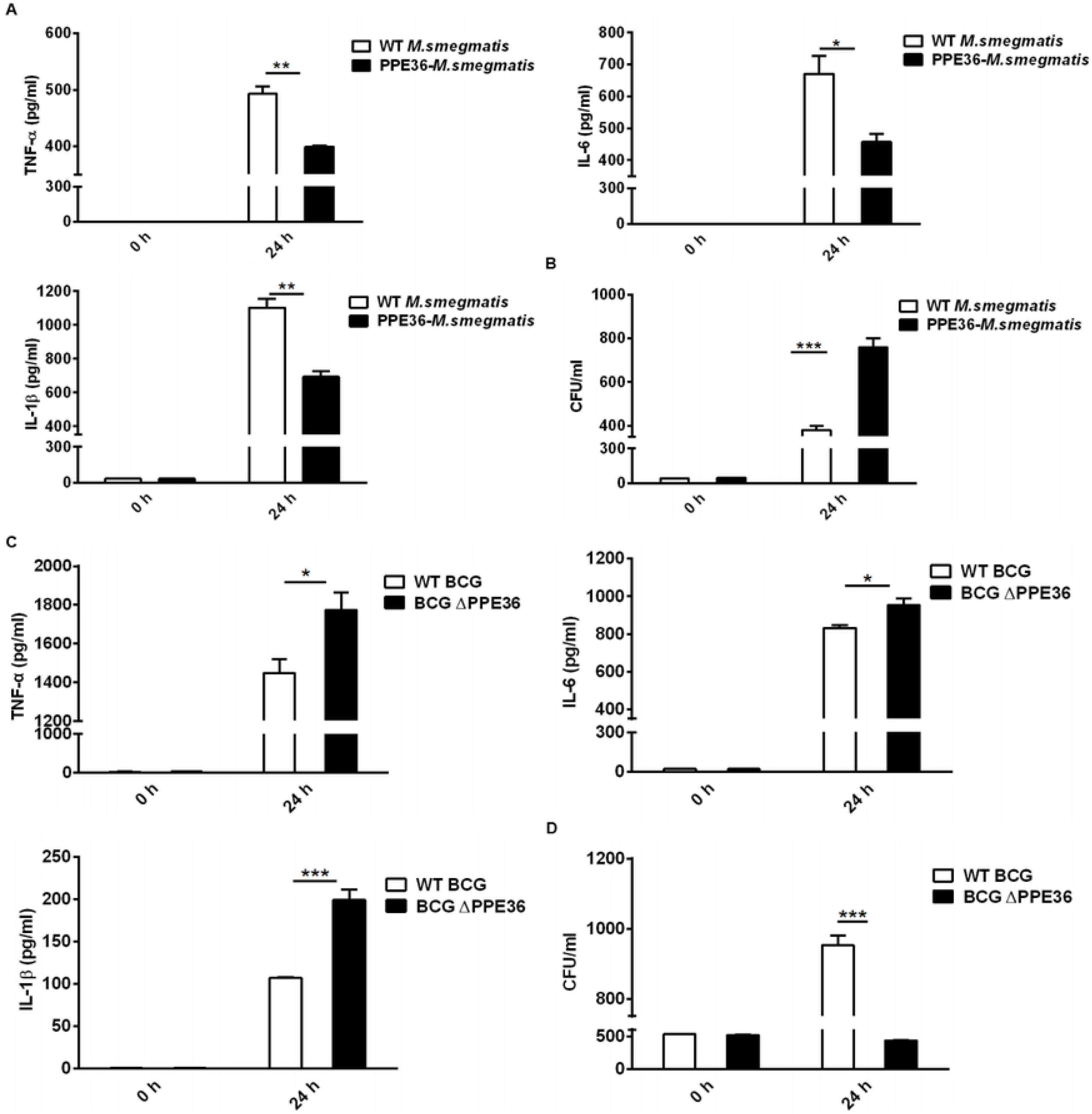
PPE36 suppressed macrophage inflammatory cytokine production. Levels of TNF-α, IL-6 and IL-1β in PPE36-M. *smegmatis* infected BMDMs. Bacterial loads in PPE36-M.*smegmatis* infected BMDMs. (C) Levels of TNF-α, IL-6 and IL-1β in BCG ΔPPE36 infected BMDMs. (D) Bacterial loads in BCG ΔPPE36 infected BMDMs. *P < 0.05, **P < 0.01, ***P <0.001.

Considering the amino acid motif of BCG-derived PPE36 is identical to that of Mtb PPE36, herein we applied PPE36-depleted BCG (BCG ΔPPE36) to explore the potential impact of PPE36 on the macrophage inflammatory responses. As shown in Figs 2C and 2D, a dramatically increased production of inflammatory cytokines (P<0.05) as well as a robustly lower intracellular bacterial load (P<0.05) were evidenced in the BCG ΔPPE36 infected cells compared with those of the WT BCG infected cells. These data indicated that PPE36 could facilitate intracellular mycobacterium survival by inhibiting inflammatory cytokine production.

### PPE36 depletion led to the increased inflammation and decreased bacterial loads in the lung tissues of mycobacterium infected mice

Inflammation plays complicated roles in the tuberculosis pathological process. An appropriate inflammatory response is essential for controlling or eliminating mycobacterium infection, while a massive response exacerbates lung tissue damage caused by the infection. Therefore, the suppressing effect of PPE36 on the mycobacterium-induced inflammation needs further verification in vivo.

Mice were intranasally infected with WT BCG or BCG ΔPPE36 for various periods, and the levels of lung inflammatory cytokines were monitored. As shown in Fig 3A, PPE 36 depletion appeared to increase the lung IL-1 β level at as early as at day 7 post infection, and this trend was even more pronounced on day 14, when lung IL-6 and TNF-α levels also began increasing, and this obvious gap maintained until day 28 post infection. These data were further supported by the decreased lung bacterial loads (Fig 3B) and the improved lung pathological observations (Figs 3C and 3D) in the BCG ΔPPE36 infected group compared with those in the control WT BCG group.

**Fig 3.**
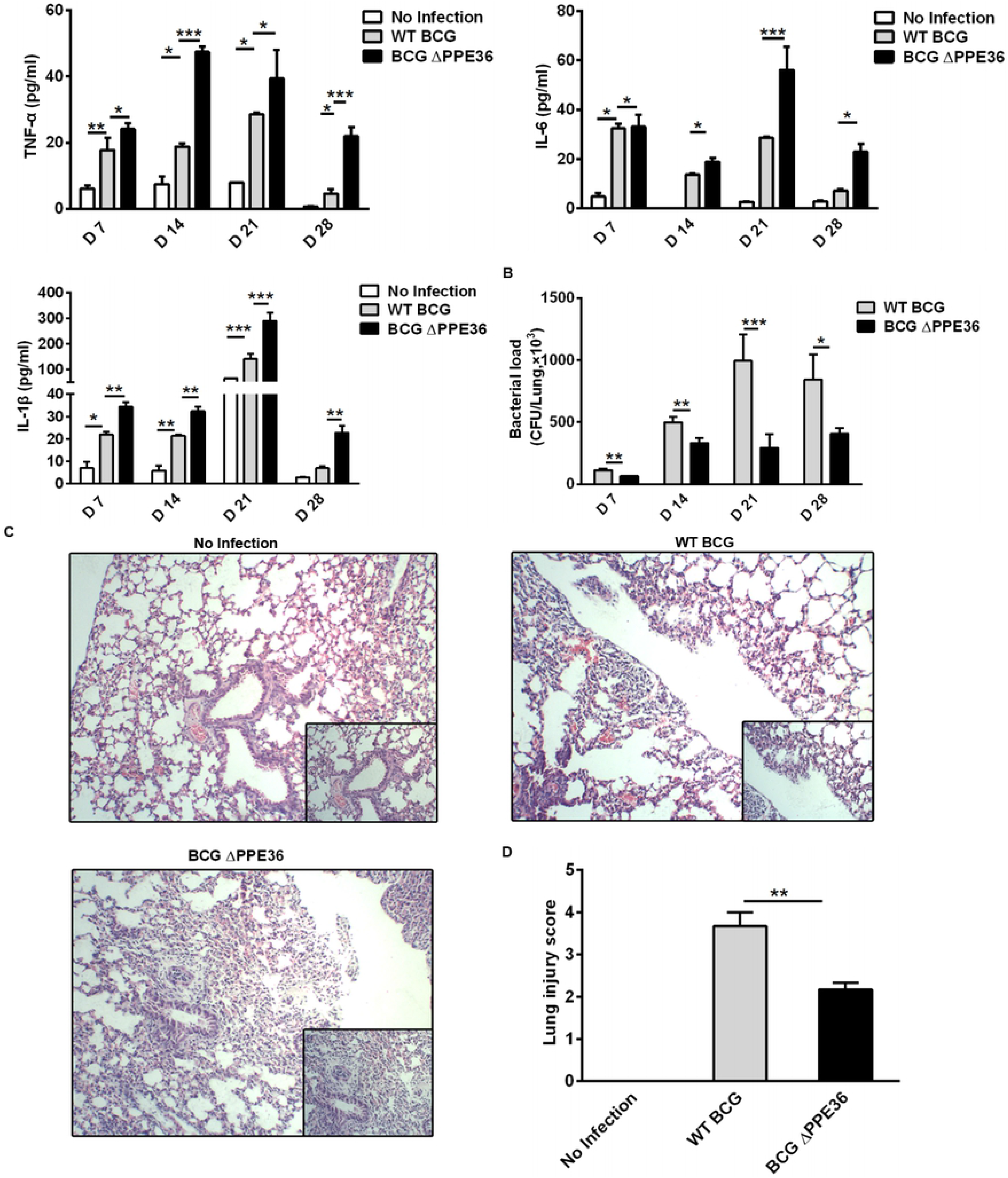
PPE36 depletion led to the increased inflammation and decreased bacterial loads in the lungs of BCG infected mice. (A) Levels of inflammatory cytokines in lung tissues of mice infected with WT BCG or BCG ΔPPE36. (B) Bacterial loads in the lung tissues of infected mice. (C) Pathological observation of lung tissues of infected mice. (D) Lung injury scores of infected mice at day 28 post infection. N.D: not detected. *P < 0.05, **P < 0.01, ***P <0.001

### Suppressive effect of PPE36 on macrophage inflammation relied on dampening the activation of NF-κB/ MAKP pathways via promoting MyD88 degradation

To explore the modulation mechanisms of PPE36 on macrophage inflammation, we evaluated PPE36 influence on the NF-κB and MAPK inflammatory pathways using a dual luciferase reporter system, which contains a pNF-κB-Luc or pAP1-Luc vector and an internal control Renilla pRL-TK luciferase vector. As shown in Figs 4A and 4B, PPE36 overexpression could robustly reduce the NF-κB activity stimulated by TNF-α as well as AP-1 activity stimulated by RacL61, the declines were as high as about 85% and 60% respectively. These data were further confirmed by the western blot results, in which PPE36 depletion led to the more potent activation of NF-κB and MAPK pathways in the BCG-infected macrophages (Figs 4C-F). Taken together, above evidence displayed that PPE36 simultaneously dampened NF-κB and AP-1 activation, suggesting that PPE36 might act on the common upstream molecules of the two pathways.

**Fig 4.**
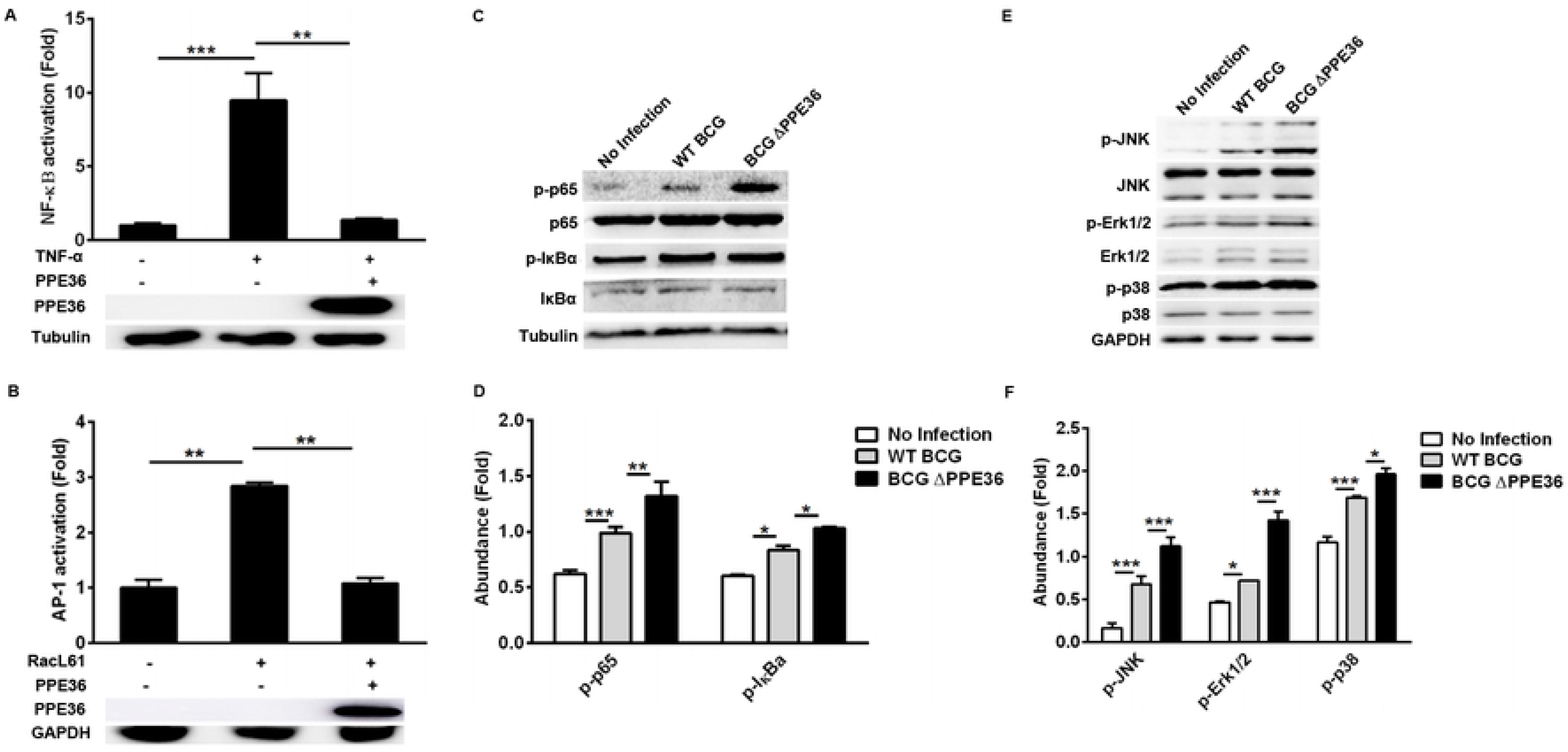
PPE36 inhibited NF-κB and MAPK activation. (A) Luciferase assays of NF-κB activation in PPE36-overexpressing 293T cells after treating with TNFα (20 ng/ml). (B) Luciferase assays of AP1 activation in PPE36-overexpressing 293T cells after transfecting pRacL61. (C-D) Western blot analysis of p-p65, p65, p-IκBα and IκBα expression in BCG or BCG∆PPE36 infected RAW264.7 cells. (E-F) Western blot analysis of p-JNK, JNK, p-Erk1/2, Erk1/2, p-p38 and p38 expression in BCG or BCG ∆PPE36 infected RAW264.7 cells. Densitometry quantification of western blot results were analyzed by Image J. *, P < 0.05, **P < 0.01, ***P <0.001.

Next, by co-transfecting plasmids encoding PPE36 and various signaling molecules (TAK1, TAB1, TAB2, TAB3, TRAF6 and MyD88), we tried to identify the potential PPE36 target by evaluating NF-κB activity with dual luciferase reporter assays. In contrast to the hardly changed NF-kB activation in cells overexpressing other signaling molecules, a robust reduced NF-κB activity was observed in the MyD88-overexpressing cells (Fig 5A), indicating that MyD88 was most likely to be the PPE36 target molecule. Consistently, we also observed a much less expression of MyD88 in the PPE36-stably expressing macrophages (Fig 5B). To further confirmed that PPE36 could reduce the MyD88 protein abundance, we transfected different doses of pPPE36-Flag plasmid into MyD88-overexpressing 293T cells, and then detected the MyD88 expression by Western Blot. As shown in Figs 5C and 5D, PPE36 potentially attenuated MyD88 expression in a dose-dependent way. All these data indicated that Mtb PPE36 could inhibit macrophage inflammation via reducing MyD88 abundance.

**Fig 5.**
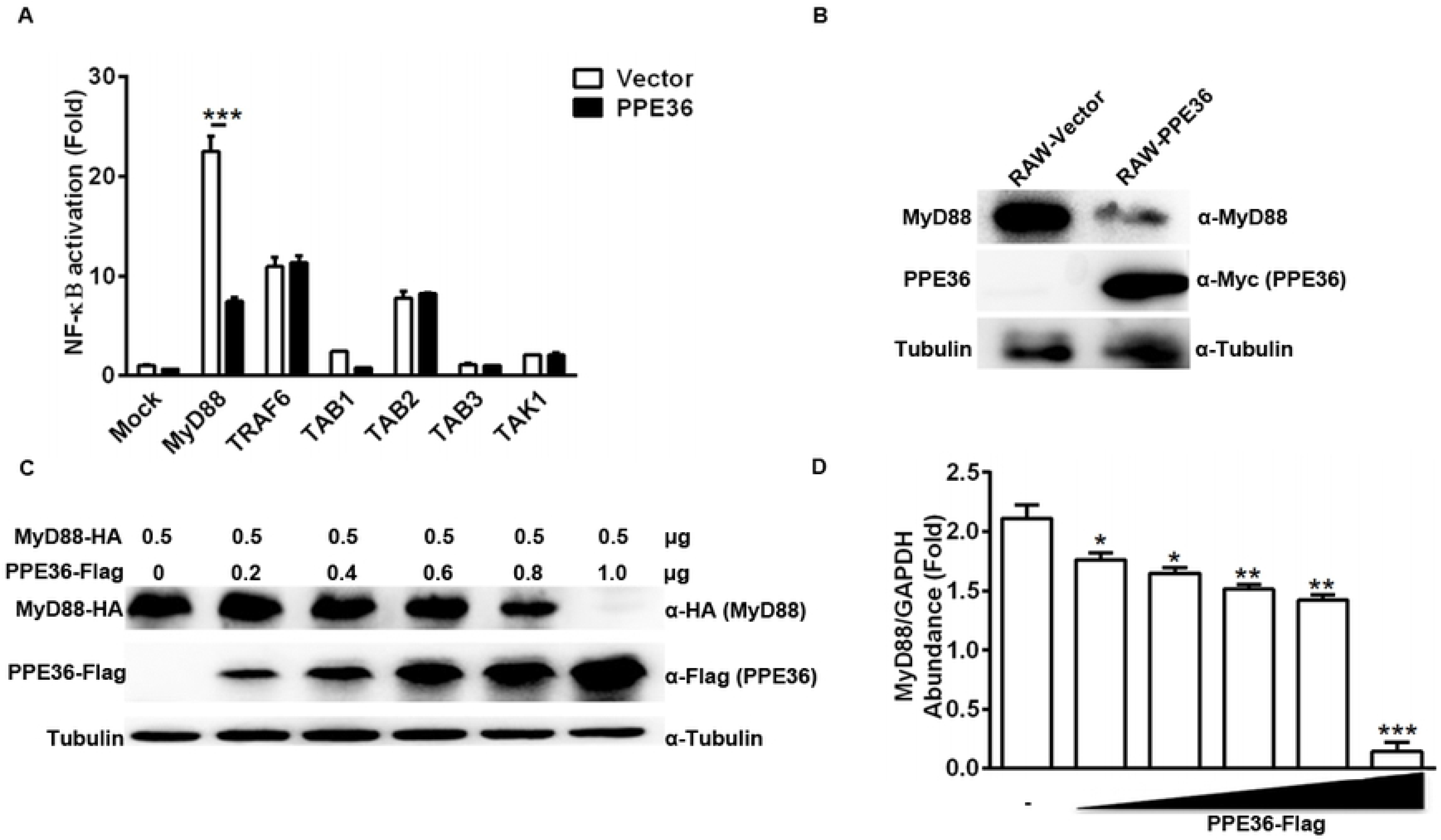
PPE36 facilitated host MyD88 protein degradation. (A) Luciferase assay of NF-κB activation in HEK293T cells co-transfected with pPPE36 and various signaling molecules (TAK1, TAB1, TAB2, TAB3, TRAF6 and MyD88). (B) MyD88 expression in PPE36-stably expressing RAW264.7 cells (RAW-PPE36). (C-D) MyD88 expression in HEK293T cell line transfected with different doses of pPPE36-Flag. *P < 0.05, **P < 0.01, ***P <0.001.

### PPE36 promoted E3 ligase Smurf1-mediated poly-ubiquitination and degradation of MyD88 protein

Protein abundance depends on the dynamic balance between its synthesis and degradation. Herein, we found that PPE36 hardly influenced MyD88 production at the transcription level (Fig 6A), suggesting that the post-translationally degradation might be changed. This deduction was further supported by the increased MyD88 abundance in PPE36-overexpressing cells in the presence of proteasome inhibitor MG132 (Fig 6B), indicating that ubiquitin (Ub)-protease degradation system was involved. In line with this, a much more intense ubiquitinated MyD88 ladder was evidenced in the PPE36-overexpressing 293T cells (Fig 6C) and macrophages (Fig 6D) compared with the control groups. These data exhibited that PPE36 could facilitate MyD88 ubiquitination and degradation.

**Fig 6.**
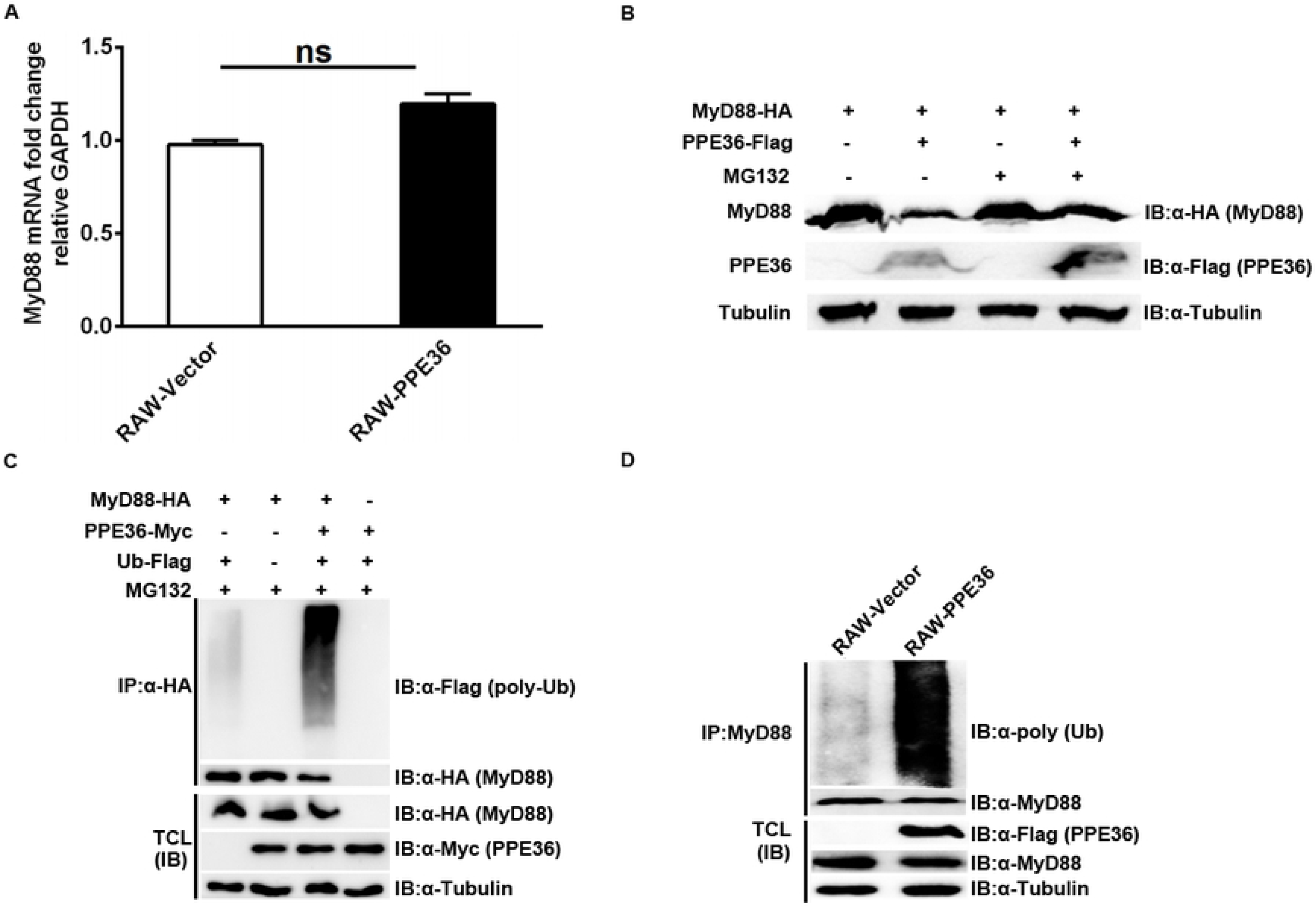
PPE36 promoted MyD88 ubiquitination. (A) MyD88 mRNA expression in PPE36-stably expressing RAW264.7 cells (RAW-PPE36). (B) MyD88 prtoein expression in HEK293T cells which were co-transfected with pMyD88-HA and pPPE36-Flag in the presence or absence of proteasome inhibitor MG132. (C) Ubiquitination level of MyD88 protein in HEK293T cells co-transfected with the indicated combinations of plasmids. (D) Ubiquitination of MyD88 protein in the PPE36-stably expressing RAW264.7 cells. TCL, total cell lysates; IP, immunoprecipitation; IB, immunoblotting.

Since no analogous functional domain of E3 ligase was predicted in PPE36 protein by searching the UbiBrowser database, we speculated that PPE36 might indirectly promote MyD88 ubiquitination through other ubiquitination ligases. To date, only E3 ligase smurf1 was shown to be involved in Mtb infection [19, 20] and participated in the MyD88 degradation, therefore we next investigated the impact of PPE36 on the interaction of MyD88 and Smurf1. We found that compared with other groups, PPE36-overexpressing group showed more Smurf1 protein binding to MyD88 (Fig 7A), indicating PPE36 obviously promoted these two protein interaction. More importantly, PPE36 efficiently favored the smurf1-mediated ubiquitination of MyD88 protein (Fig 7A). Similar increased binding of Smurf1 and MyD88 was also seen in PPE36-stably expressing macrophages (Fig 7B). Meanwhile PPE36-caused MyD88 ubiquitination was obviously abolished when Smurf1 was down-regulated (Fig 7C), indicating that Smurf1 was the primary E3 ligase in this process. These data indicated that PPE36 promoted Smurf1-mediated ubiquitination and degradation of MyD88.

**Fig 7.**
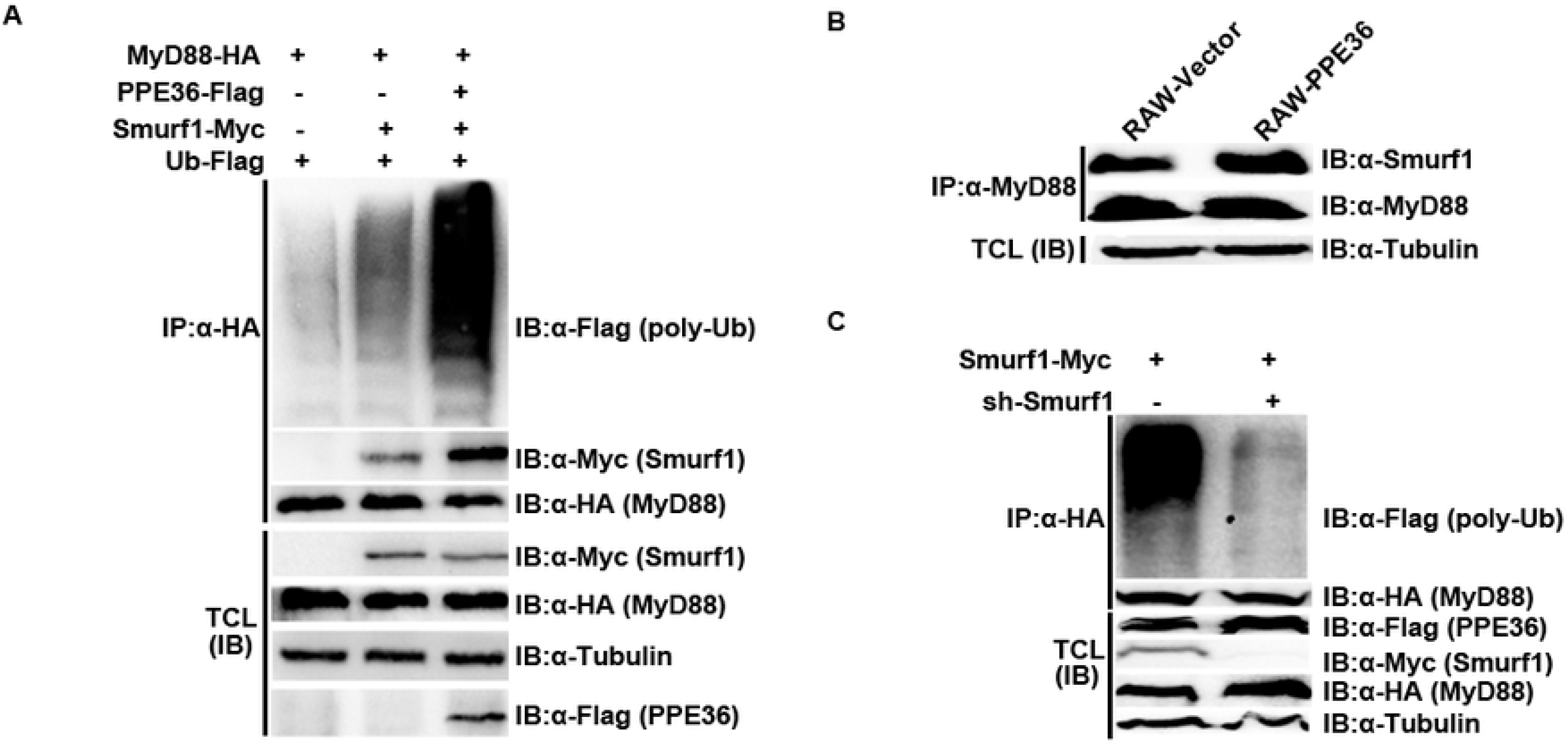
PPE36 promoted Smurf1-mediated ubiquitination and degradation of MyD88 protein. (A) Ubiquitination of MyD88 protein and the interaction of MyD88 with E3 ligase Smurf1 in HEK293T cells which were co-transfected with pPPE36-Flag, pMyD88-HA, pSmurf1-Myc as well as pUb-Flag. (B) Interaction of MyD88 with the E3 ligase Smurf1 in PPE36-stably expressing RAW264.7 cells (RAW-PPE36). (C) HEK293T cells were transfected with pSmurf1-Myc for 24h, and then transfected with sh-Smurf1 or control shRNA, 12 h later, MyD88 ubiquitination level was monitored by immunoprecipitation assays. All data were representative of at least three independent experiments. TCL, total cell lysates; IP, immunoprecipitation; IB, immunoblotting.

## Discussion

Mtb is the world’s most successful pathogen. It invades hosts mainly through respiratory tracts, and the first innate immune cells it encounters are alveolar macrophages, which armed with plenty of immune mechanisms to detect and eliminate intracellular pathogens. While after thousands of years of reciprocal evolution with the human beings, Mtb has evolved a series of complex immune escape mechanisms that can successfully evade or modulate macrophage immune responses to ensure its successful survival. Studies in murine mycobacterium infection models showed that lung bacterial loads significantly increase during the innate immunity stage, yet stopped growing and achieved the platform when adaptive immune responses initiated [21], further displayed the eximious abilities of mycobacterium to evade innate immunity.

PPE/PE are two unique families of multi-gene proteins in Mtb. They encode two major glycine-rich protein families, accounting for nearly 10% bacterial genomic encoding capability [22]. PPE/PE proteins possess a comparatively conserved N-terminal and a highly variable C-terminal, which were conducive to escape the host immune attack [5, 23]. In addition, PPE/PPE proteins were exclusively restricted to the virulent mycobacterium strains, and this phenomenon was further confirmed by this study. We showed that PPE36 expression was significantly higher in the virulent H37Rv strain and clinical isolates, compared with that in the attenuated BCG strain, suggested that PPE36 might play an essential part in the mycobacterium infection and disease.

Protein localization is extremely critical to its function. However, at present, the location and function of most PPE/PE proteins are still largely unknown. Previous studies reported that some secreted PPE proteins can interact with the host cell surface PRR or enter the host cytoplasm to regulate signaling pathways [17, 24–27]. Some studies also showed that cell-membrane or cell-wall associated PE/PPE proteins also regulated cell signaling pathways in a variety forms. For example, the cell-wall associated protein PPE32 could increase the cytokine secretion by activating endoplasmic reticulum stress [28]. PPE44 was reported to promote IL-12 and IL-6 expressions via activating NF-κB/ERK1/2/p38 pathways [29]. Interestingly, some cleaved products of surface-exposed PPE proteins also possessed the abilities to regulate intracellular signaling pathways. For example, the cleaved C terminal of PPE37 could transport to nuclear and activate the caspase 3-dependent apoptosis pathway [17]. In this study, we found that PPE36 could both inhibit NF-κB and MAPK signaling pathways. Correspondingly, BCG ΔPPE36 infection led to the increased inflammatory cytokines (TNF-α, IL-6 and IL-1β) and the reduced lung bacteria loads. We further identified that MyD88 protein was the important target of Mtb PPE36, as the inflammation-suppressive effect was dampened when MyD88 was deficient.

Actually, MyD88 plays an indispensable role during Mtb infection. It was reported that IFN-γ-induced full-scale macrophage activation can only be carried out when MyD88 was present [30]. Compared with the wide type counterparts, MyD88-deficient mice were more sensitive to Mtb infection, and more easily progressing to granulomas with massive inflammation and necrosis [31, 32]. Our previous study also reported another tuberculosis protein EspR, which could bind to MyD88 and inhibit the TLR pathway activation as well as subsequent macrophage inflammation [33]. In line with our data, Bandyopadhyay U and colleagues [34] showed that Mtb protein sulfotransferase Rv3529c (Stf1) inhibited TLR2-mediated immune response by disrupting the interaction of MyD88 with IRAK1. Of interest, TcpC protein of virulent Escherichia coli also exhibited the ability to regulate the TLR4/NF-κB signaling pathway by acting on MyD88 [35]. It seemed that MyD88 might act as a common and key target for diverse bacterial immune-modulating proteins. Herein, we found that PPE36 significantly reduced the intracellular MyD88 expression via promoting its ubiquitination and proteasomal degradation, without influencing its gene transcription.

So far there is no report showing any PPE molecule functioning as a ubiquitin E3 ligase, although their activities as lipase or hydrolase enzyme have been reported [36]. Additionally, we did not find out any functional E3 ligase domain in PPE36 with the help of UbiBrowse database previously reported [37], therefore we speculated that PPE36 may indirectly regulate MyD88 ubiquitination through other E3 ligases.Several E3 ubiquitination ligases have been reported to be involved in MyD88 ubiquitination and degradation, like Smurf1 [20, 38], Nrdp1 [39] and Cbl-b [40]. Although all these molecules have been reported to regulate macrophage inflammatory response, only Smurf1 was shown to be involved in Mtb infection. Therefore, we focused on investigating whether PPE36 could affect the interaction of MyD88 and Smurf1. We found that in the presence of PPE36, the binding of Smurf1 and MyD88 was significantly increased, and the subsequent MyD88 ubiquitination level was also obviously increased. While when down-regulating the Smurf1 expression, the ability of PPE36 to promote MyD88 ubiquitination was robustly dampened, suggesting that PPE36 mainly promoted the ubiquitination and degradation of MyD88 through E3 ubiquitin enzyme Smurf1. Of note, Smurf1 has been reported as an important factor for restricting Mtb replication in macrophages, helping hosts withstand pulmonary tuberculosis in mice, and controlling mycobacterium growth in human macrophages [20].These were mainly achieved by Smurf1-promoted autophagy [20]. Ironically, Mtb PPE36 in turn exploits this host defense molecule smurf1 to inhibits host inflammation by promoting MyD88 degradation, further demonstrating the unparalleled adaptability of Mtb to its living environment and powerful gene regulation ability.

Taken together, our data demonstrated the new role of Mtb PPE36 in regulating the host immune response, and showed that PPE36 could promote the Smurf1-mediated MyD88 ubiquitination and degradation, and then dampen host innate immunity and facilitate Mtb survival.

## Materials and methods

### Mycobacteria strains

*M. smegmatis*, *M. bovis* BCG and *M. tuberculosis* (Mtb) H37Rv strains were grown at 37°C on Middlebrook 7H10 agar (BD Diagnostic Systems) supplemented with 10% ADC, or in Middlebrook 7H9 broth medium (BD Diagnostic Systems) supplemented with 10% OADC, 0.5% glycerol and 0.05% Tween-80. Herein, *M. smegmatis* strains included WT *M.smegmatis* (WT-*M.s megmatis*), PPE36-overexpressing *M.smegmatis* (PPE36-*M. smegmatis*); *M. bovis* BCG strains included WT BCG, PPE36-deleted BCG (BCG ∆PPE36). Both BCG and *M.smegmatis* were obtained from Shanghai Institutes for Biological Sciences and Shanghai Institutes for Biological Sciences. H37Rv strain was obtained from Affiliated Hospital of Zunyi Medical University. BCG ΔPPE36 strain was purchased from Gene Optimal Inc (Shanghai, China). PPE36-*M.smegmatis* was prepared as transfecting the plasmid pMV261-PPE36 into *M.smegmatis* by electroporation according to standard procedures [41]. Mtb clinical isolates were collected from the sputum and bronchoalveolar lavage fluids (BALF) of tuberculosis patients from January 2008 to December 2018 in the Tuberculosis Division of Respiratory and Critical Care Medicine at Affiliated Hospital of Zunyi Medical University, Guizhou, China. The obtained clinical specimens were processed following the protocols recommended by WHO [42]. Isolates were cultured on Löwenstein– Jensen solid slants, and grown colonies were further identified to the species level using 2-thiophene carboxylic acid and para-nitrobenzoic acid selective media.

### Cells, plasmids and antibodies

HEK293T cells and murine macrophage cell line RAW264.7 were cultured in DMEM medium (Hyclone) supplemented with 10% Fetal Bovine Serum (Hycolne), 0.1 mg/ml streptomycin, 100 U/ml penicillin at 37 °C with 5% CO_2_. Bone marrow derived macrophages (BMDMs) were prepared as previously described [33].

The mammalian expression plasmids pPPE36-Flag, pPPE36-Myc, pMyD88-HA were constructed in our laboratory; plasmids pRacL61, pTRAF6, pTAB1, pTAB2, pTAB3, pTAK1 were provided by Cuihua Liu (Chinese Academy of Sciences, China); plasmids pSmurf1-Myc, pUbiquitin (Ub)-Flag as well as Smurf shRNA (sh-Smurf1) were provided by Hui Zheng (Soochow University, China). The dual luciferase reporter assay vectors pNF-kB-luc, pAP1-luc, pRL-TK were purchased from Clontech. The mycobacterium shuttle vector pMV261 (provided by Dr. Honghai Wang,Fudan University, China) was used to overexpress Mtb PPE36 in *M. smegmatis.*

Primary antibodies against GAPDH, p-p65, p65, p-IκBα, IκBα, Tubulin, p-JNK, JNK, p-Erk1/2, p-Erk1/2, p-p38, p38 (Cell Signaling Technology) were used for western blot assays. Antibodies against MyD88, HA, Myc, Ubiquitin, Smurf1 (Cell Signaling Technology) and Flag (Sigma) were used for immunoprecipitation assays. HRP-conjugated anti-mouse or anti-rabbit secondary antibodies were purchased from Southern-Biotech.

### Quantification of PPE36 expression by real-time PCR

The total RNAs were extracted from various mycobacteria and transcribed into cDNAs as previously described [43], and PPE36 expressions were quantified by absolute quantification real-time PCR using the standard curve method [44]. Standard curves were generated using a dilution series of 1010 to 103 copies per microliter of the linearizing pPPE36-Flag as a template. Using the average molecular weight of the product and Avogadro's constant, the number of copies per unit volume was calculated as described previously [44–46].

### Genexpert Assay determining bacterial load

GeneXpert assays were used to determine bacterial loads in the clinical sputum and BALF samples as described [47]. In brief, 1ml sputum or 200 μl BALF decontaminated samples were added into 2 ml of sample reagent and transferred into a test cartridge. The cartridge then was inserted into the test platform of GeneXpert instrument. Threshold Cycle (Ct) values for each of five rpoB probes were recorded and used to estimate bacterial load. A Ct value of 40 was assigned if GeneXpert was negative for Mtb detection.

### Mycobacterium infection

Murine macrophage cell line RAW264.7 or BMDMs were infected with *M. smegmatis,* PPE36-*M.smegmatis,* WT BCG or BCG ΔPPE36 at a multiplicity of infection (MOI) of 10, and then cells were harvested at different time points.

Six to eight week-old C57BL/6 mice were purchased from Shanghai SLAC Laboratory Animal Co., Ltd. (Shanghai, China), and were housed using standard humane animal husbandry protocols at Soochow University. All animal experiments were approved by the Institutional Animal Care and Use Committees of Soochow University. Mice were intranasally infected with 1 × 107 colony forming units (CFU) of WT BCG or BCG ΔPPE36, and sacrificed at 7, 14, 21 or 28 days.

### Colony-forming unit (CFU) counting

CFU counting was applied to determine the bacterial loads of infected macrophages and mouse lung tissues. Aliquots of cell lysates were serially diluted 10-fold with water plus 0.05%Triton X-100. Homogenized infected lung tissues were diluted with PBS. And 50 μl of cell lysates or tissue homogenates were added to 7H10 plates and cultured for 3 weeks, and then CFU counting was performed.

### Lung injury evaluation

Lung tissues were fixed with 4% (v/v) paraformaldehyde, embedded in paraffin for slicing, and then subjected to hematoxylin and eosin staining. The sections were analyzed by lung injury score to score lung inflammation and damage [48]. Microscopic scoring criteria of lung injury were graded from 0 to 4: Grade 0: normal lung morphology; Grade 1: mild intra-alveolar edema and mild inflammatory cell infiltration; Grade 2: moderate intra-alveolar edema and moderate inflammatory cell infiltration; Grade 3: severe alveolar edema, severe inflammatory cell infiltration, and focal hemorrhage; and Grade 4: disseminated inflammatory cell infiltration and destruction in alveolar structure.

### Luciferase reporter assays

For detecting NF-κB activity detection, HEK293T cells were co-transfected with pNF-κB-luc (1 μg) and pRL-TK (50 ng) plasmids in the presence of pPPE36 (1 μg) or control plasmid (1 μg) for 24 h using Lipofectamine 2000 reagent (Invitrogen). And then cells were treated with TNF-α (20 ng/ml) for 12 h. For detecting JNK and p38 activation, HEK293T cells were co-transfected with pRacL61(1 μg), pAP1-luc (1 μg) and pRL-TK (50 ng) in the presence of PPE36-Flag (1 μg) or control plasmid (1 μg) for 36 h. Cells were lysed and subjected to luciferase activity detection according to manufacturer instructions (Promega). All reactions were performed in triplicate.

### Western blot analysis

Cells lysates were fractionated by 10% SDS-PAGE, transferred onto PVDF membranes and incubated with primary antibodies against tubulin, GAPDH, PPE36 or molecules associated with NF-κB and AP1 signaling pathways. After washing with PBST three times, membranes were further incubated with appropriate HRP-conjugated anti-rabbit or anti-mouse secondary antibodies. Signals were detected with enhanced chemiluminescence (ECL) by Amersham Imager 600(GE), and quantified with ImageJ.

### Immunoprecipitation and ubiquination assays

HEK293T cells were co-transfected with pMyD88-HA (0.5 μg), pPPE36-Flag (1.5 μg), pSmurf1-Myc (1.5 μg) and pUb-Flag (50 ng) for 36 h, and then cell lysates were prepared and immunopricipitated with anti-HA or anti-MyD88 beads. The precipitates were then immunoblotted with anti-Ub, anti-Smurf1, anti-Myc, anti-HA, anti-Flag or anti-MyD88 antibodies.

### Statistical analysis

All data were expressed as mean±SEM. Statistical differences in 2 groups or more than 2 groups were respectively assessed by Student’s t test or One-way ANOVA followed by Bonferroni test using GraphPad Prism version 6.0 (GraphPad Software Incorporated). P values less than 0.05 were considered to be significant.

## Acknowledgments

We thank Cuihua Liu from Chinese Academy of Sciences, for providing the plasmids of pRacL61, pTRAF6, pTAB1, pTAB2, pTAB3, pTAK1. We thank Hui Zheng from Institute of Biology and Medical Sciences for providing the plasmids of pSmurf1-Myc, pUbiquitin (Ub)-Flag as well as Smurf1 shRNA (sh-Smurf1). We thank Lin Chen from the Affiliated Hospital of Zunyi Medical University for providing the clinical isolate strains.

